# High WEE1 expression is independently linked to poor survival in multiple myeloma

**DOI:** 10.1101/2024.09.20.613788

**Authors:** Anish K. Simhal, Ross Firestone, Jung Hun Oh, Viswatej Avutu, Larry Norton, Malin Hultcrantz, Saad Z. Usmani, Kylee H. Maclachlan, Joseph O. Deasy

## Abstract

Current prognostic scores in multiple myeloma (MM) currently rely on disease burden and a limited set of genomic alterations. Some studies have suggested gene expression panels may predict clinical outcomes, but none are presently utilized in clinical practice. We therefore analyzed the MMRF CoMMpass dataset (N=659) and identified a high-risk group (top tertile) and a low-risk group (bottom tertile) based on WEE1 expression sorted in descending order. The tyrosine kinase WEE1 is a critical cell cycle regulator during the S-phase and G2M-checkpoint. Abnormal WEE1 expression has been implicated in multiple cancers including breast, ovarian, and gastric cancers, but has not until this time been implicated in MM. PFS was significantly different (p <1e-9) between the groups, which was validated in two independent microarray gene expression profiling (GEP) datasets from the Total Therapy 2 (N=341) and 3 (N=214) trials. Our results show WEE1 expression is prognostic independent of known biomarkers, differentiates outcomes associated with known markers, is upregulated independently of its interacting neighbors, and is associated with dysregulated P53 pathways. This suggests that WEE1 expression levels may have clinical utility in prognosticating outcomes in newly diagnosed MM and may support the application of WEE1 inhibitors to MM preclinical models. Determining the causes of abnormal WEE1 expression may uncover novel therapeutic pathways.

## Introduction

Multiple myeloma (MM) is a hematologic malignancy associated with a malignant proliferation of plasma cells [1]. Although the disease is usually responsive to upfront therapies, MM remains incurable even in patients who achieve undetectable levels of disease, with relapse considered largely inevitable [2]. The genomic makeup of MM is highly heterogeneous, and different studies have identified multiple subtypes associated with varying prognostic outcomes using different data modalities [1,3–5]. Standard methods to prognosticate the length of progression-free survival (PFS) include the International Staging System (ISS) [6], Revised ISS (R-ISS) [7], and the Second Revision of the ISS (R2-ISS) [8]. These tools rely on surrogates for disease burden and identification of specific tumor cytogenetic abnormalities. These scoring systems each have a PFS concordance index (c-index) below 60%, leaving room for improvement [9,10].

In addition to providing genomic information, scoring systems informed by gene expression have been proposed for prognostication, including GEP70 and SKY92 [11,12]. These expression-based signatures have shown potentially complementary information to ISS staging [13]. In [5], we conducted a large unsupervised genomic network study where we applied a novel measure of network connectivity, Ollivier-Ricci curvature (ORC), to RNA-sequencing (RNA-seq) and copy number alteration (CNA) data from newly diagnosed MM (NDMM) patients. We examined patterns of gene-gene interactions in MM and identified novel pathways and genes associated with poor prognosis. By examining the impact of gene expression via a network, we identified a novel eight-gene signature: *BUB1, MCM6, NOSTRIN, PAM, RNF115, SNCAIP, SPRR2A*, and *WEE1*. Of these eight genes, *WEE1* was the only gene that was included in a previously published gene signature, GEP70 [14]. Furthermore, *WEE1* was the most prognostic for PFS, suggesting it might play a role in MM. However, the role of the *WEE1* in MM has not been thoroughly studied, and much remains unknown about its prognostic significance with respect to known biomarkers of MM.

*WEE1* is a tyrosine kinase involved in multiple aspects of the cell cycle process, including the G1-S checkpoint, S phase, and G2-M checkpoint [15,16], but believed to exert its most significant clinical impact in the G2-M checkpoint. For non-cancerous cells, DNA damage is often repaired at the G1-S checkpoint. In cancerous cells, the G1-S checkpoint may be deficient, and therefore, cancerous cells rely on the G2-M checkpoint for DNA damage repair [17]. In the G2-M checkpoint, *WEE1* regulates cyclin-dependent kinase 1 (*CDK1*) [18–20], with high *WEE1* expression suppressing *CDK1* expression and maintaining the cell in a DNA repair state [21,22]. Conversely, low *WEE1* expression correlates with a rise in *CDK1* expression, which allows the cell to enter mitosis [18].

*WEE1* inhibition has been shown to dysregulate the cellular machinery associated with the first stage of mitosis in the G1-S transition [23], and can induce apoptosis by forcing mitotic entry [24]. For a cell to successfully complete the cell cycle, *WEE1* expression levels must rise and fall in relation to each stage of the cycle. High *WEE1* expression has recently been shown to be associated with disease aggressiveness in some solid tumors including breast cancer [25], ovarian cancer [26], and melanoma [27–30]. Several *WEE1* inhibitors are currently in phase 2 clinical trials; these trials are evaluating the therapeutic efficacy of *WEE1* inhibition [31,32]. *WEE1* inhibitors have also shown promise in other cancer types including sarcomas [33] and breast cancers [34], as well as hematological malignancies [35].

In MM, preclinical studies have shown promising results when inhibiting *WEE1* in cell lines and mouse models in conjunction with other factors [36–40]. *WEE1* inhibitors, in combination with bortezomib, can induce apoptosis in MM cell lines more efficiently than bortezomib alone [36,37]. further, in [38], the authors show that bortezomib in combination with a DNA damage response (DDR) inhibitor targeting *ATM*/*ATR*/*WEE1* triggers apoptosis. In [39], the authors examine the relationship between *WEE1* and *CHK1* in MM, and report that targeting both kinases induces apoptosis in MM cell lines. In [40], the authors suggest targeting *CTPS1* in conjunction with either *CHEK1, ATR*, or *WEE1* inhibition can induce apoptosis in MM cell lines.

In this study, we show that high *WEE1* expression defines a high-risk subtype of MM, independent of both known markers of MM and treatment types. *WEE1* expression has comparable prognostic value as compared to the traditional MM ISS. Additionally, high *WEE1* expression is not reflected by corresponding changes in expression throughout the transcriptome. The high *WEE1* expression subtype is characterized by dysregulation of the P53 pathway. Together, this work suggests that in a subpopulation of MM patients, WEE1 may play an outsized role and should be studied as a potential therapeutic target.

## Methods

In this study, we applied a variety of bioinformatic and machine learning-based methods to MM datasets to examine the role of *WEE1* in MM.

### CoMMpass data

The RNA-seq and copy number alterations (CNA) data used is from the Multiple Myeloma Research Foundation’s CoMMpass dataset, release version 19. Further information on the data collection and curation methods has previously been published [41,42]. The details of the patients selected for this study along with the preprocessing and feature computations are described in detail in [5]. Briefly, for inclusion in this study, subjects must have RNA-Seq and CNA data extracted from the bone marrow plasma cells before the start of treatment and both demographic and survival information available (N=659). Gene inclusion was based on overlap with the Human Protein Reference Database (HPRD) [43].

### Gene expression profiling (GEP) data

The GEP data used is from the University of Arkansas’s Total Therapy 2 (TT2, N=341) and Total Therapy 3 (TT3, N=214) trials. The details of these trials are described in [44,45]. Briefly, the plasma cells were collected via a bone marrow biopsy of newly diagnosed MM patients before treatment and gene expression profiling data was collected. TT2 & TT3 were different treatment regimens. Note that for this dataset, event-free survival (EFS) was reported.

### High-risk group membership

For each data modality — RNA-seq and GEP — patients’ WEE1 expression values were sorted in descending order and the top tertile was labeled as *WEE1*-high and the bottom tertile was labeled as *WEE1*-low. The center third was not considered in this study.

### Prognosis and confounder analysis

The prognosis was modeled using Kaplan Meier (KM) survival curves for progression-free survival (PFS). To determine the effect of WEE1 relative to known biomarkers of MM, we used a multivariate Cox proportional hazards model [46] with the RNA-Seq data to predict PFS. In it, we modeled nine markers: hyper APOBEC, chromothripsis, hyperdiploidy, *MAF* translocation, *MYC* translocation, t(4;14), t(11;14), TP53 mutation, and gain 1q21. As outlined in [5], hyperdiploidy was defined by more than 2 gains involving >60% of the chromosome affecting chromosomes 3, 5, 7, 9, 11, 15, 19, or 21. Mutational signatures were assessed using *mmsig* (https://github.com/UM-Myeloma-Genomics/mmsig), a fitting algorithm designed for MM to estimate the contribution of each mutational signature in each sample [47]. APOBEC-mutational activity was calculated by combining *SBS2* and *SBS13*, with the top 10% being defined as hyper-APOBEC [48,49]. The complex structural variant chromothripsis was defined by manual curation according to previously published criteria [50]. High-risk and low-risk groups were analyzed separately to see which factors differed between the groups. To show the prognostic effect of *WEE1*, irrespective of known biomarkers, KM survival curves for PFS stratified by each factor were plotted.

### Machine learning analysis

We used random survival forests [51] to determine the prognostic value of *WEE1*, its gene network neighbors, and ISS. Briefly, random survival forests offer the advantages of random forests with the addition of incorporating survival information including event duration and censorship information. *WEE1* neighbors were extracted from the STRING database [52]. *WEE1* neighbors were defined as genes which have a known interaction with *WEE1* with a probability greater than 0.7. The neighboring genes were considered to see if changes in *WEE1* expression were reflected by changes in expression of known interacting genes. ISS staging was provided by the CoMMpass dataset. We used the concordance index (c-index) as the evaluation metric. *WEE1* expression was predicted using random forest regression models to see if neighboring genes contained signal relevant to the abnormal increase in *WEE1* expression. Feature importances were computed using the permutation importance method in sci-kit-learn and the fifteen most importances are reported [53]. The full parameter details of the models used are available on GitHub (www.github.com/aksimhal/WEE1-in-MM). Models were evaluated using five-fold cross-validation repeated ten times.

### Differential gene expression analysis

To see differences in patterns of gene expression between the *WEE1*-high and *WEE1*-low cohorts, we computed the differential gene expression using DESeq2 [54]. The p-values from this analysis were corrected for multiple hypothesis testing using BH-FDR method. Genes with a corrected p-value less than 0.05 and an absolute log2 fold change greater than two were considered significant. To see which pathways become dysregulated in *WEE1*-high, we used the Gene Set Enrichment Analysis tool to evaluate the selected genes [55,56]. The utilized pathways are from the hallmark gene set collection from the human molecular signatures database (MSigDB) [57].

### Data and code availability

The code and instructions for how to use them are available for download at www.github.com/aksimhal/WEE1-myeloma. The Multiple Myeloma Research Foundation’s CoMMpass data is available for download at www.research.mmrf.org. TT2 and TT3 are available at GSE24080.

## Results

### Data overview

Genomic and clinical characterization of MM outcomes were stratified by *WEE1* expression using the CoMMpass dataset (N=659). The mean age was 62.5 ± 10.7 years and 60% were male; ISS distribution was 35/35/30%, and 53% received an autologous stem cell transplant (ASCT). An overview of the differences between the *WEE1*-high and *WEE1*-low groups is provided in Table 1. While some of the known markers of MM are significant between the two groups, including age, hyperdiploidy, t(11;14), *MAF* and *MYC* translocations, chromothripsis, hyper APOBEC, gain 1q21, and TP53 mutational status, ISS is not. For the validation datasets, TT2 and TT3, baseline clinical data and gene expression data were available. For TT2, the mean age was 56.3 ± 9.8 years and 57% male; for TT3, the mean age was 58.6 ± 8.8 years and 67% male.

**Table 1.** Difference in CoMMpass data patient characteristics between WEE1-high and WEE1-low cohorts. The majority of MM markers differ significantly between the two groups; however, ISS does not. Key: for gain 1q21, 0 = diploid, 1 = gain (3 copies), 2 = amplification (4 or more copies). For TP53 aberration, 0 = diploid, 1 = either deletion or mutation, 2 = biallelic loss. Certain markers not available for all subjects. BTZ: Bortezomib, CFZ: Carfilzomib.

|  | WEE1-low (N=218) | WEE1-high (N=224) | FDR p-value |
| --- | --- | --- | --- |
| **Age** | **63.4** | **61.1** | **3.31E-02** |
| Sex | M: 114; F: 104 | M: 139; F: 85 | 5.22E-02 |
| ISS | (I): 75; (II): 88; (III): 48 | (1): 79; (2): 73; (3): 69 | 8.77E-02 |
| Treatment | combined BTZ/IMiDs-based: 124; BTZ-based: 38; combined IMiDs/CFZ-based: 21; IMiDs-based: 14; CFZ-based: 11; combined BTZ/IMiDs/CFZ-based: 9; combined BTZ/CFZ-based: 1 | combined BTZ/IMiDs-based: 89; combined IMiDs/CFZ-based: 50; BTZ-based: 45; CFZ-based: 20; IMiDs-based: 11; combined BTZ/IMiDs/CFZ-based: 9 | - |
| Hyperdiploidy | 151/189 | 58/170 | 9.53E-18 |
| t(4;14) | 22/204 | 20/181 | 1.00E+00 |
| t(11;14) | 11/204 | 69/181 | 4.30E-15 |
| MAF translocation | 4/204 | 22/181 | 1.48E-04 |
| MYC translocation | 39/204 | 19/181 | 3.31E-02 |
| Chromothripsis | 42/204 | 56/181 | 2.57E-02 |
| Hyper APOBEC | 6/204 | 25/179 | 2.04E-04 |
| Gain 1q21 | (0): 141; (1): 47; (2): 1 | (0): 105; (1): 46; (2): 19 | 1.39E-04 |
| TP53 aberration | (0): 157; (1): 15; (2): 0 | (0): 114; (1): 22; (2): 13 | 1.48E-04 |

### WEE1 is prognostic for outcomes in RNA-seq and GEP datasets

In the RNA-seq data from the CoMMpass dataset, differences in PFS between *WEE1*-high and *WEE1*-low cohorts are statistically significant (p <1e-9), as shown in Figure 1A. These results are validated in the TT2 and TT3 datasets (Figures 1B, 1C). Note this effect is not observed in the CNA data from the CoMMpass dataset.

**Figure 1.**
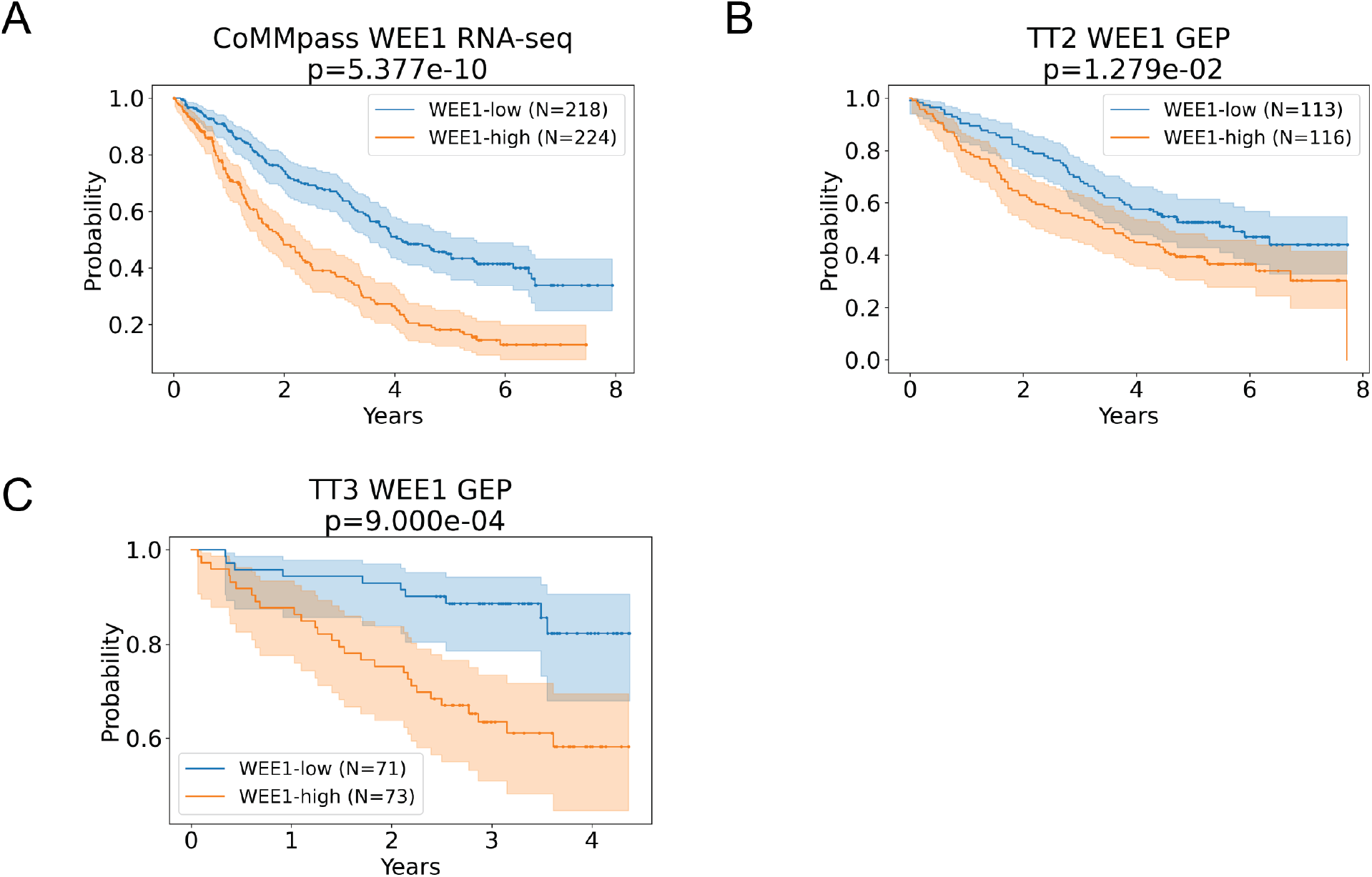
Prognostic value of WEE1 expression from RNA-seq and GEP data. A) Progression free survival (PFS) based on CoMMpass RNA-seq data showing the two-year difference in median PFS with a p-value of less than 1e-9. B & C) Event free survival of the Total Therapy 2 and Total Therapy 3 cohorts gene expression profiling (GEP) data, respectively, showing diverging outcomes with a P<0.05.

### Multivariate modeling shows that WEE1 is an independent prognostic factor in MM

Multivariate Cox proportional hazards modeling shows that the prognostic effect of *WEE1* is independent of known MM markers, including those shown to be significant in Table 1. The prognostic effect is independent of hyperdiploidy, t(4;14), t(11,14), *TP53* status, as well as emerging risk factors, the complex structural variant chromothripsis and APOBEC mutational activity, shown in Figure 2A and Supplemental Table 1A. When examining only the WEE1-high cohort, none of the markers significantly predicted PFS (Figure 2C, Supplemental Table 1B). Similarly, in the *WEE1*-low, none of the markers significantly predicted PFS (Figure 2B, Supplemental Table 1C).

**Figure 2.**
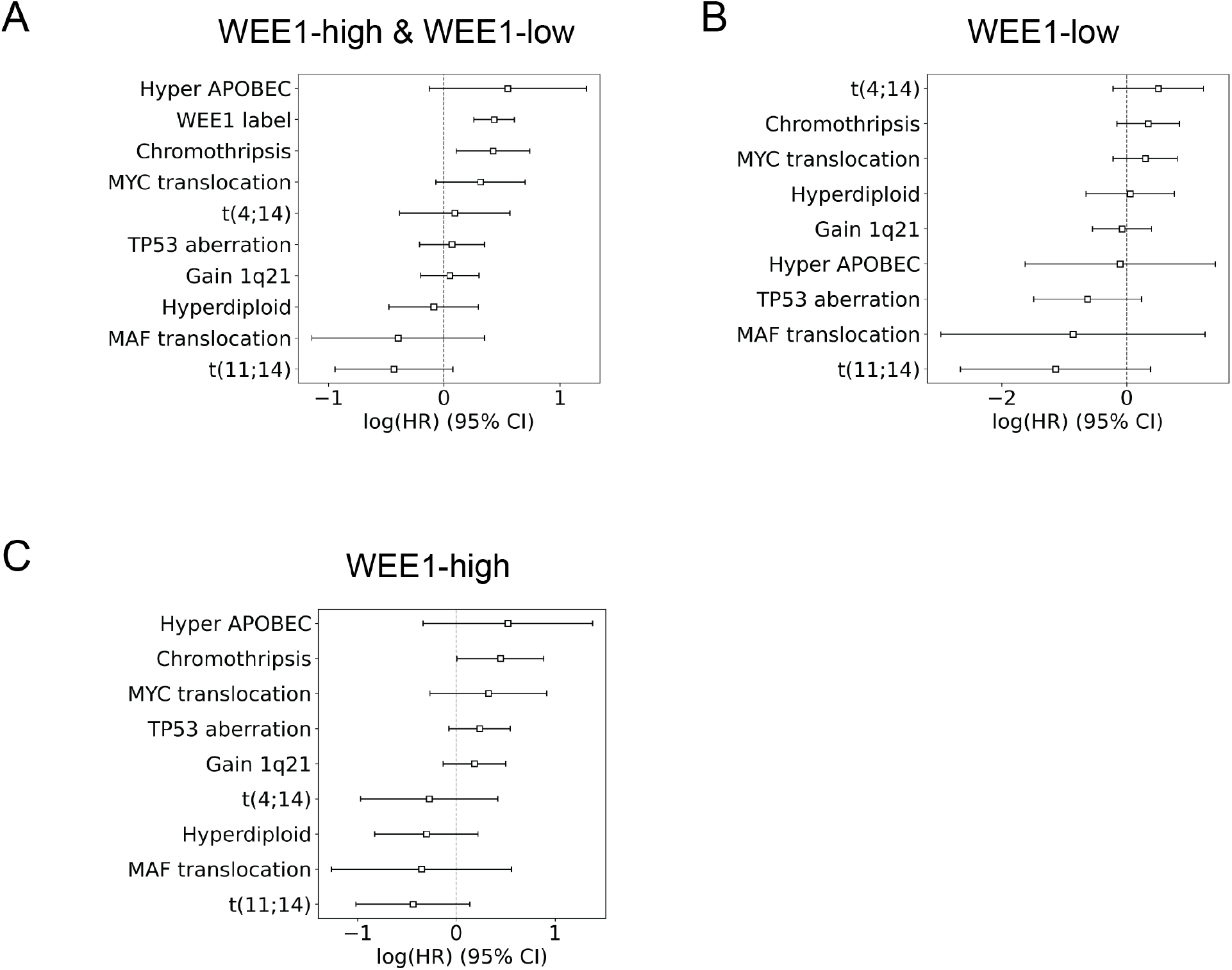
Cox proportional hazards (CPH) modeling of MM markers and *WEE1* expression. A) Coefficients of the multivariate CPH model show *WEE1* to be the most significant prognosticator. B & C) Within the WEE1-high and WEE1-low cohorts, none of the markers are significant for PFS after FDR-BH correction. TP53 aberration status — 0 = diploid, 1 = either deletion or mutation, 2 = biallelic loss. Certain markers not available for all subjects.

### WEE1 is prognostic for outcomes independent of known biomarkers

The *WEE1*-high and *WEE1*-low cohorts have statistically significantly different PFS outcomes when stratifying for each known MM marker. KM plots show significant separation when looking at groups defined by the presence of hyperdiploidy, t(11;14), *MAF & MYC* translocations, chromothripsis, and *TP53* deletion (Figure 3, Supplemental Figure 1). KM plots were also significant when looking at the groups defined by the lack of a known MM marker (Figure 3, Supplemental Figure 2). *WEE1* cohort membership differentiates outcomes by an average of 1.98 years in cohorts with a marker, and 2.18 years in cohorts without the marker (Table 2).

**Table 2.** The difference in median progression free survival (PFS) is based on a given biomarker. “Positive” indicates the cohort which has the listed feature. “Negative” indicates the cohort which does not have the listed feature. The difference is calculated as the median PFS of the WEE1-low group minus the median PFS of the WEE1-high group. ND is defined as “no data” and indicates that the LR group did not reach the median PFS mark.

| Feature name | WEE1-high<br>PFS (years) | WEE1-low<br>PFS (years) |
| --- | --- | --- |
| Hyperdiploidy | 1.838 | 2.685 |
| t(4;14) | 1.115 | 2.436 |
| t(11;14) | ND | 1.956 |
| MAF translocation | ND | 1.921 |
| MYC translocation | 2.121 | 2.436 |
| Chromothripsis | 2.427 | 2.427 |
| Hyper APOBEC | 2.378 | 1.942 |
| TP53 deletion | ND | 1.608 |

**Figure 3.**
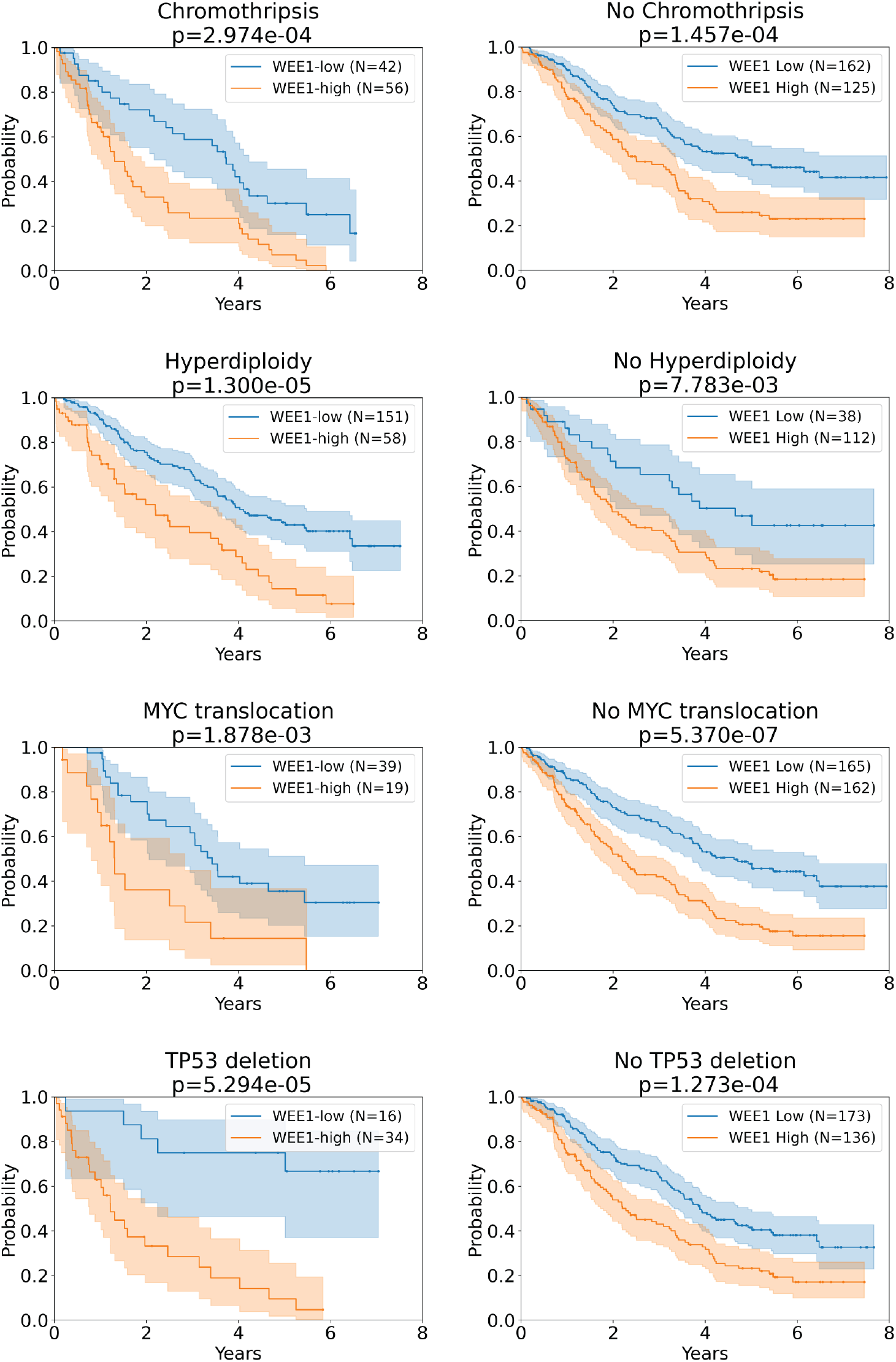
Kaplan Meyer curves stratified by MM markers show the prognostic signal in *WEE1* expression. *WEE1* expression defines prognosis regardless of marker type. The top row represents the cohort with a given feature, and the bottom row represents the cohort without the given feature. In both cases, *WEE1* defined low-risk and high-risk groups as separate outcomes with a median PFS difference of two years.

### WEE1 is prognostic for outcomes independent of treatment type

The *WEE1*-high and *WEE1*-low cohorts have statistically significantly different PFS outcomes when stratifying the treatment options listed in the CoMMpass dataset. Autologous stem cell transplant (ASCT), bortezomib/immunomodulatory agents (IMIDs), bortezomib, and carfilzomib/IMIDs cohorts were all significantly different when stratified by *WEE1*-high and *WEE1*-low (Figure 4). The mean difference in PFS is 1.91 years.

**Figure 4.**
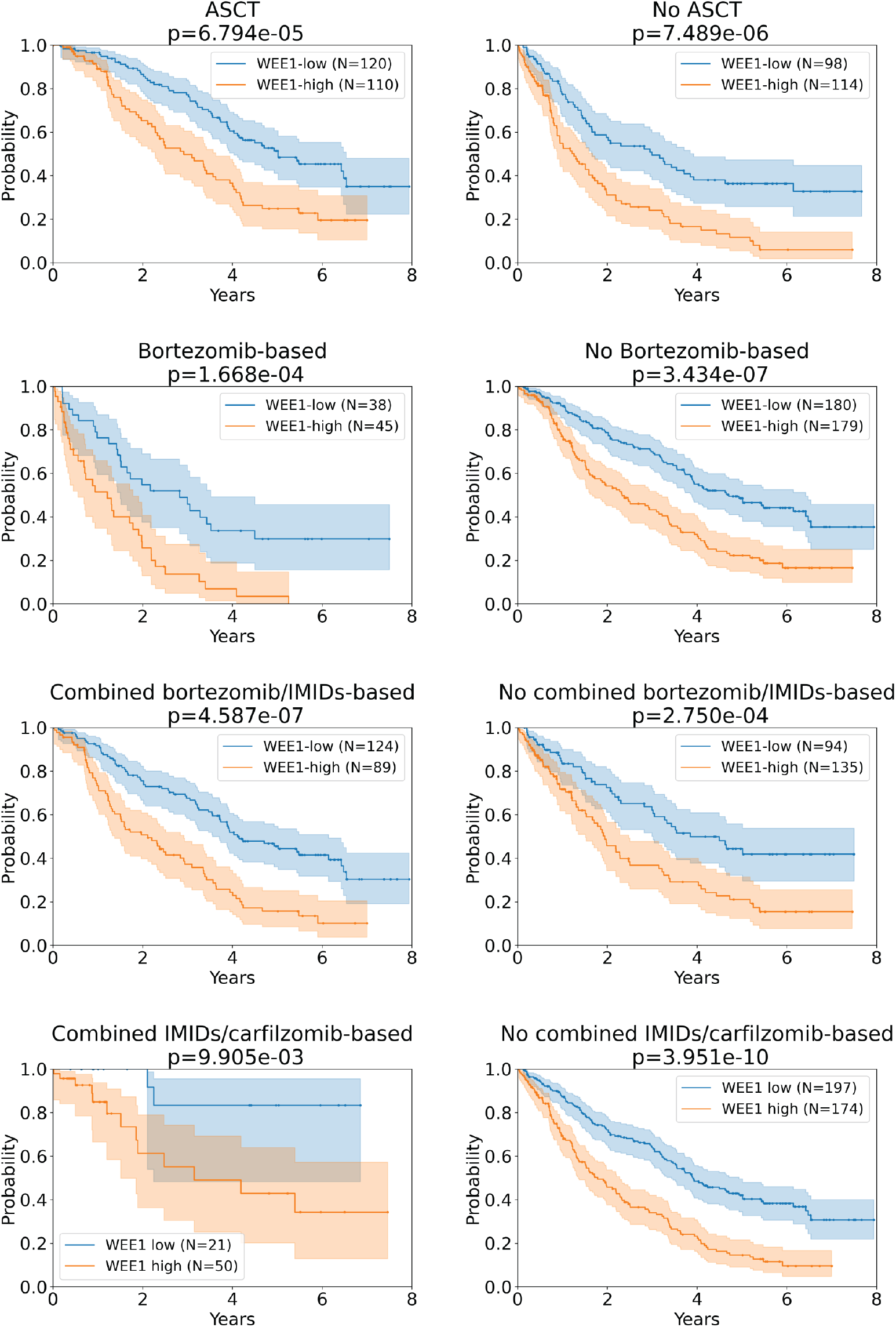
Kaplan Meyer curves show the effect of *WEE1* expression on treatment type. The top row is the cohort that received a treatment type, and the bottom row is the cohort that did not receive the treatment type.

### WEE1 expression has comparable prognostic value as ISS

RNA-seq based *WEE1* expression has comparable *prognostic* value (c-index: 0.58 ± 0.04) as ISS (c-index: 0.61 ±0.03). Combining *WEE1* and ISS has a c-index of 0.63±0.03.

### WEE1-high cohort is 3.2x less predictable than the WEE1-low cohort

As *WEE1* expression increases, the relationship between *WEE1* and genes known to interact with *WEE1* becomes dysregulated. When modeling *WEE1* expression with known interacting genes, the prediction error increases by 3.2 times between the *WEE1*-high and *WEE1*-low cohorts. In the *WEE1*-low cohort, the known interacting genes that contribute more than 5% to the prediction are *CDK1, CHEK1, CDT1, AURKB*, and *PLK1* (Figure 5A). In the *WEE1*-high cohort, the genes are *CDC25B, HSP90AA1, CDK6, PLK1, CDR2, SKP2*, and *CDK2* (Figure 5B).

**Figure 5.**
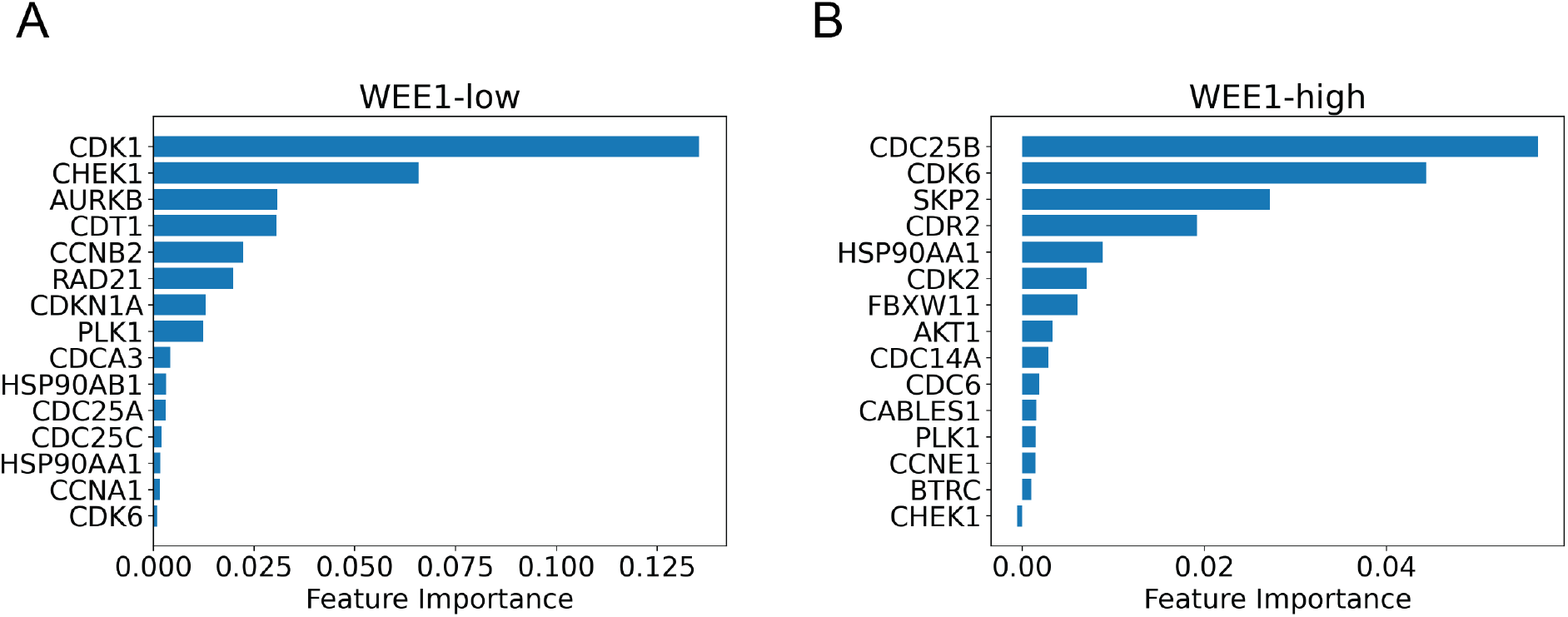
Random forest feature importance plots. RF modeling of WEE1 expression in the WEE1-high cohort is 3.2x more inaccurate than WEE1 expression modeling in the WEE1-low cohort. A) Feature importance plot showing the informative features for predicting *WEE1* RNA-seq in the *WEE1*-low group. B) Feature importance plot showing the informative features for predicting *WEE1* RNA-seq in the *WEE1*-high group.

### P53 pathway-related genes are differentially expressed between WEE1-high & WEE1-low cohorts

A differential gene expression analysis between the *WEE1*-high and *WEE1*-low groups identified 146 overexpressed genes and five underexpressed genes. Overexpressed genes are part of three pathways: P53, downregulated UV response, and mitotic spindle. Only five genes were under-expressed: *FPR1, IFNA5, LRP2, POU2F3*, and *RAB11FIP1*.

## Discussion

Prognostic markers in MM rely on either assessment of tumor burden or specific cytogenetic abnormalities; transcriptional characteristics of myeloma are not currently considered in this setting. Here, we have identified that high *WEE1* expression represents an independent biomarker prognostic of poor outcomes in newly diagnosed MM, and that this effect is independent of known cytogenetic risk factors and treatment strategies (Figures 1-3). This includes the common metric of staging — ISS. Random survival forest modeling showed that *WEE1* expression alone has as much prognostic power as ISS staging. These findings were seen both retrospectively using the CoMMpass dataset and independently validated in two prospective MM data sets. Differential gene expression analysis showed that the P53 pathway is the most significantly affected pathway in the *WEE1*-high cohort.

Random forest modeling of the local *WEE1* genomic network showed that the overexpression of *WEE1* is not correlated with an increase or decrease in any genes locally connected with *WEE1*. Increased *WEE1* expression was not reflected in a rise in the expression of any other cell cycle kinases, such as *PLK1* or *CDK1*. Random forest modeling of the low-risk group showed an association with *CDK1*, which follows known biology. In our defined high-risk group, *CDK1* was not in the top 15 genes most associated with the high-risk *WEE1* signal. In this group, *CDC25B*, a phosphatase-encoding gene, replaces *CDK1* as the most influential marker. This further suggests that *WEE1* expression represents an independent prognostic marker that is likely not merely reporting on another known cytogenetic risk factor.

*WEE1* is a key player during the cell cycle, and its specific roles in the S phase and the G2M checkpoint are well documented. *WEE1* acts as a tumor suppressor gene in certain types of breast cancer. However, for the majority of solid and blood cancers, such as ovarian cancer and acute lymphoblastic leukemia, *WEE1* acts as an oncogene. Further work is needed to understand the role of increases in *WEE1* expression in MM as these findings can enable new *WEE1* directed treatments in MM patients with MM and other malignancies.

Of note, differences in PFS among patients with *TP53* deletions when stratifying by *WEE1* expression were remarkably large. Patients with *TP53* deletions often have the poorest clinical outcomes with MM treatment across multiple published datasets. Additionally, differential gene expression analysis between the high-risk and low-risk groups showed that genes associated with the hallmark P53 pathway were differentially expressed. *TP53* regulates DNA damage in the G1-S checkpoint. Faulty P53 function may lead to a larger reliance on *WEE1* activity to maintain genomic integrity. If both *TP53* and *WEE1* are abnormal, it is possible that DNA repair becomes dysfunctional.

We have demonstrated that stratification of MM patients with *TP53* deletions by MM cell *WEE1* expression may represent an alternative method of risk stratifying patients. Additionally, our data suggests that *WEE1* inhibition may be especially effective in patients with altered P53 pathways, though further investigation is needed to identify if the observed association is causal. There are currently five *WEE1* inhibitors in clinical trials [58] for other cancer types which will advance our understanding of the efficacy of *WEE1* inhibition, the exact mechanism of its actions, as well as a possible new treatment option for MM patients. Further investigation into the apparent centrality of WEE1 in predicting outcomes, especially at a biological level, is required to validate it as a critical biomarker.

## Acknowledgments

This study was supported in part by an MSK Cancer Center Support grant (P30 CA008748), The Simons Foundation, and a Breast Cancer Research Foundation grant (BCRF-17-193).

## Competing Interests

SZU: Research funding: Amgen, BMS/Celgene, GSK, Janssen, Merck, Pharmacyclics, Sanofi, Seattle Genetics, Takeda. Consulting/Advisory Board: Abbvie, Amgen, BMS, Celgene, Genentech, Gilead, GSK, Janssen, Sanofi, Seattle Genetics, SecuraBio, SkylineDX, Takeda, TeneoBio.

## Notes

### Competing Interest Statement

The authors have declared no competing interest.

