## Supplemental Figures for "High WEE1 expression is independently linked to poor survival in multiple myeloma"

**Supplemental tables and figures for "High WEE1 expression is independently linked to poor survival in multiple myeloma"**

Anish K. Simhal*[1], Ross Firestone[2], Jung Hun Oh[1], Viswatej Avutu[3], Larry Norton[3], Malin Hultcrantz[2], Saad Z. Usmani[2], Kylee H. Maclachlan[2], Joseph O. Deasy[1]

[1]Department of Medical Physics, Memorial Sloan Kettering Cancer Center, New York, United States of America, [2]Myeloma Service, Department of Medicine, Memorial Sloan Kettering Cancer Center, New York, United States of America, [3]Department of Medicine, Memorial Sloan Kettering Cancer Center, New York, United States of America

**Supplemental Table 1. Multivariate CPH modeling results.** A) Multivariate CPH modeling results for PFS show WEE1 HR membership to be the most predictive marker, followed by chromothripsis. This implies that the other eight markers do not have an effect larger than WEE1 HR membership. B) Within the LR group, none of the markers are significant. C) Within the HR group, none of the markers are significant.

**A**

| Covariate | Coefficient | SE | Lower 95% | Upper 95% | p |
|---|---|---|---|---|---|
| WEE1 label | 0.433 | 0.089 | 0.258 | 0.607 | 0.000 |
| Hyperdiploid | -0.091 | 0.198 | -0.478 | 0.297 | 0.647 |
| t(4;14) | 0.092 | 0.244 | -0.386 | 0.570 | 0.706 |
| t(11;14) | -0.433 | 0.260 | -0.943 | 0.077 | 0.096 |
| MAF translocation | -0.397 | 0.381 | -1.143 | 0.349 | 0.297 |
| MYC translocation | 0.314 | 0.196 | -0.070 | 0.698 | 0.109 |
| Chromothripsis | 0.422 | 0.162 | 0.105 | 0.738 | 0.009 |
| Hyper APOBEC | 0.549 | 0.346 | -0.129 | 1.228 | 0.113 |
| TP53 code | 0.067 | 0.144 | -0.214 | 0.349 | 0.639 |
| gain1q21 | 0.049 | 0.129 | -0.204 | 0.303 | 0.704 |

**B**

| Covariate | Coefficient | SE | Lower 95% | Upper 95% | p |
|---|---|---|---|---|---|
| Hyperdiploid | 0.056 | 0.360 | -0.650 | 0.761 | 0.877 |
| t(4;14) | 0.504 | 0.368 | -0.217 | 1.225 | 0.170 |
| t(11;14) | -1.140 | 0.774 | -2.657 | 0.377 | 0.141 |
| MAF translocation | -0.861 | 1.077 | -2.971 | 1.249 | 0.424 |
| MYC translocation | 0.296 | 0.261 | -0.216 | 0.808 | 0.257 |
| Chromothripsis | 0.342 | 0.254 | -0.156 | 0.839 | 0.178 |
| Hyper APOBEC | -0.106 | 0.774 | -1.623 | 1.410 | 0.891 |
| TP53 code | -0.627 | 0.438 | -1.486 | 0.231 | 0.152 |
| gain1q21 | -0.076 | 0.241 | -0.548 | 0.395 | 0.751 |

**C**

| Covariate | Coefficient | SE | Lower 95% | Upper 95% | p |
|---|---|---|---|---|---|
| Hyperdiploid | -0.305 | 0.268 | -0.829 | 0.220 | 0.255 |
| t(4;14) | -0.276 | 0.355 | -0.971 | 0.420 | 0.437 |
| t(11;14) | -0.441 | 0.295 | -1.020 | 0.137 | 0.135 |
| MAF translocation | -0.356 | 0.465 | -1.268 | 0.557 | 0.445 |
| MYC translocation | 0.324 | 0.302 | -0.268 | 0.916 | 0.284 |
| Chromothripsis | 0.445 | 0.225 | 0.005 | 0.886 | 0.047 |
| Hyper APOBEC | 0.522 | 0.439 | -0.339 | 1.383 | 0.235 |
| TP53 code | 0.235 | 0.159 | -0.076 | 0.546 | 0.138 |
| gain1q21 | 0.182 | 0.162 | -0.135 | 0.500 | 0.260 |

**Supplemental Figure 1. Kaplan Meyer curves stratified by MM markers show the prognostic signal in *WEE1* expression.** *WEE1* expression defines prognosis regardless of marker type. The top row represents the cohort with a given feature, and the bottom row represents the cohort without the given feature. In both cases, *WEE1* defined low-risk and high-risk groups as separate outcomes with a median PFS difference of two years.

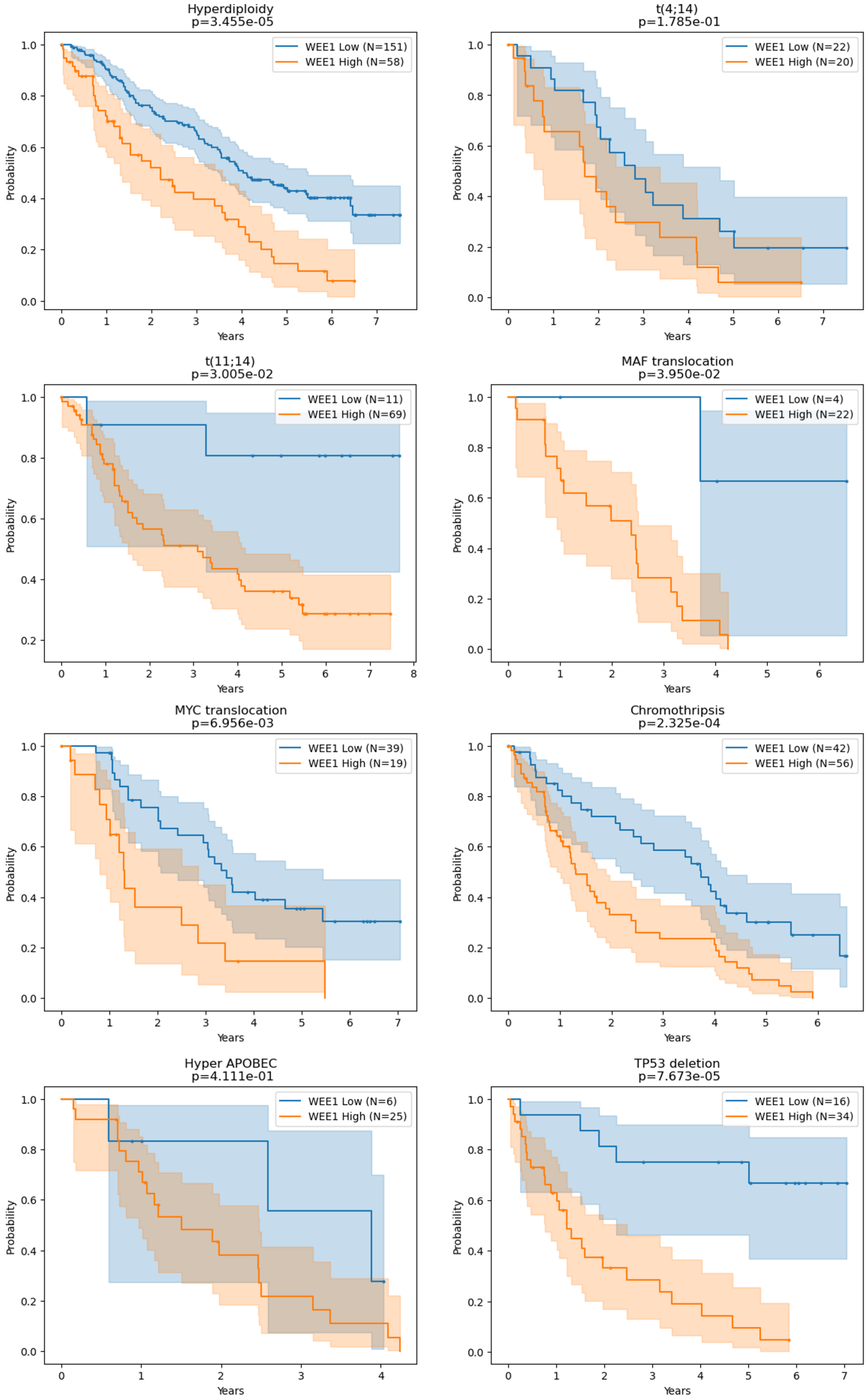
Hyperdiploidy
p=3.455e-05
WEE1 Low (N=151)
WEE1 High (N=58)
t(4;14)
p=1.785e-01
WEE1 Low (N=22)
WEE1 High (N=20)
t(11;14)
p=3.005e-02
WEE1 Low (N=11)
WEE1 High (N=69)
MAF translocation
p=3.950e-02
WEE1 Low (N=4)
WEE1 High (N=22)
MYC translocation
p=6.956e-03
WEE1 Low (N=39)
WEE1 High (N=19)
Chromothripsis
p=2.325e-04
WEE1 Low (N=42)
WEE1 High (N=56)
Hyper APOBEC
p=4.111e-01
WEE1 Low (N=6)
WEE1 High (N=25)
TP53 deletion
p=7.673e-05
WEE1 Low (N=16)
WEE1 High (N=34)
Probability
Years

**Supplemental Figure 2. Kaplan Meyer curves stratified by lack of MM markers show the prognostic signal in *WEE1* expression.** *WEE1* expression defines prognosis regardless of marker type. The top row represents the cohort with a given feature, and the bottom row represents the cohort without the given feature. In both cases, *WEE1* defined low-risk and high-risk groups as separate outcomes with a median PFS difference of two years.

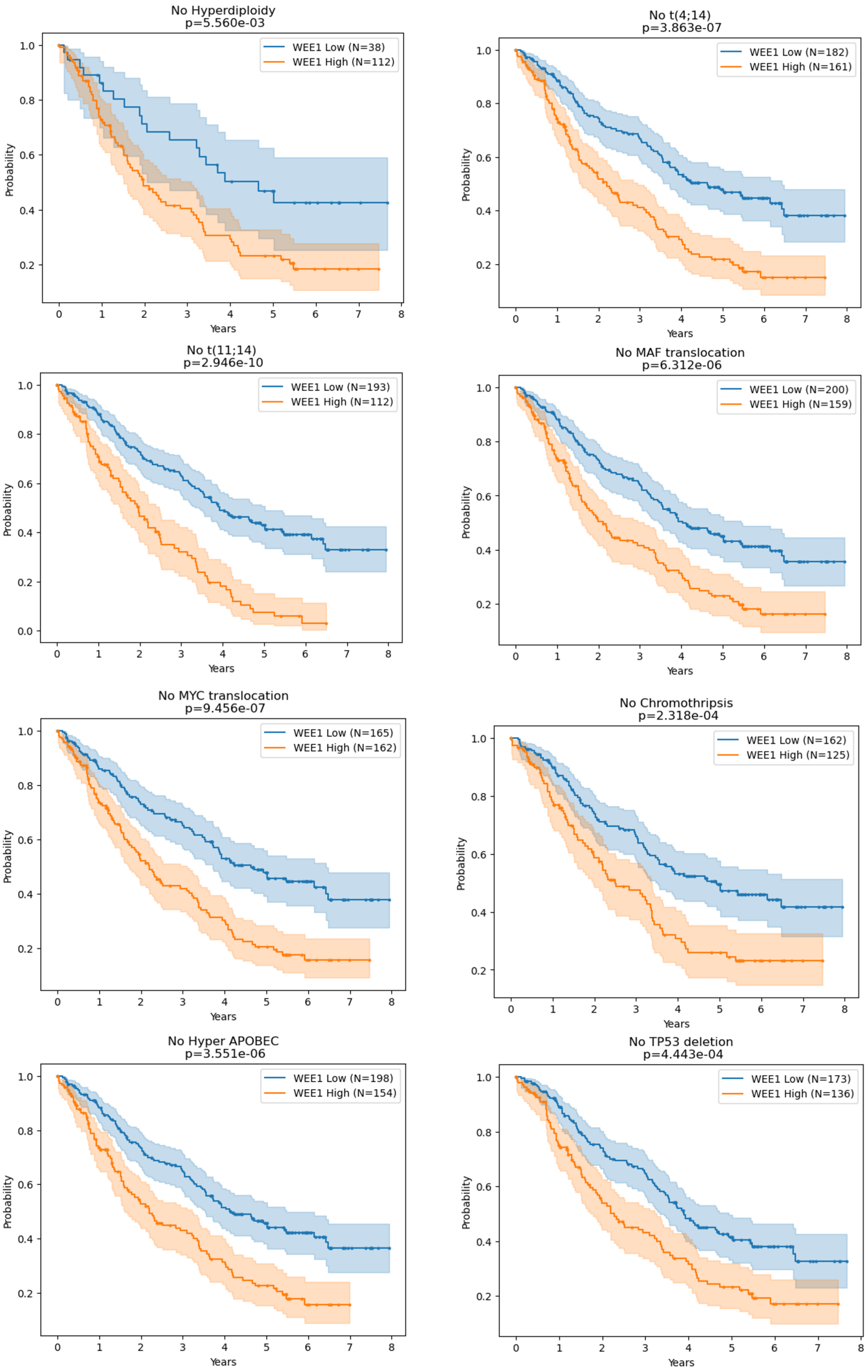
No Hyperdiploidy
p=5.560e-03
WEE1 Low (N=38)
WEE1 High (N=112)
No t(4;14)
p=3.863e-07
WEE1 Low (N=182)
WEE1 High (N=161)
No t(11;14)
p=2.946e-10
WEE1 Low (N=193)
WEE1 High (N=112)
No MAF translocation
p=6.312e-06
WEE1 Low (N=200)
WEE1 High (N=159)
No MYC translocation
p=9.456e-07
WEE1 Low (N=165)
WEE1 High (N=162)
No Chromothripsis
p=2.318e-04
WEE1 Low (N=162)
WEE1 High (N=125)
No Hyper APOBEC
p=3.551e-06
WEE1 Low (N=198)
WEE1 High (N=154)
No TP53 deletion
p=4.443e-04
WEE1 Low (N=173)
WEE1 High (N=136)
Probability
Years
